# ProFinder: Candidate constitutive promoter prediction across the bacterial tree for genetic tool development in non-model species

**DOI:** 10.64898/2026.09.23.753901

**Authors:** Anupam Gautam, James W. Marsh

## Abstract

Reliable constitutive promoters for non-model bacteria are lacking, resulting in a bottleneck for genetic tool development. Existing computational promoter predictors are trained on model Gammaproteobacteria and degrade on phylogenetically distant or compositionally atypical genomes. Here we present ProFinder, a computational method that integrates three biologically informed criteria for promoter prediction: a conserved 98-gene constitutive marker panel, an empirically derived intergenic search window, and a 72-motif σ^70^ motif catalogue built from diverse lineages. From an input bacterial assembly, ProFinder returns a ranked shortlist of putative constitutive promoters. It matches the precision of full-genome classifiers and is more robust to GC extremes. Applied to 56,742 bacterial genomes spanning 149 phyla, ProFinder returned candidate promoters for 99.76%, delivering a pan-bacterial catalogue of pre-computed constitutive promoter parts to accelerate chassis development across the bacterial tree. ProFinder can be accessed at: https://plabase.cs.uni-tuebingen.de/profinder/.

## Introduction

Promoters are the principal drivers of transcription and serve as the fundamental genetic switch that controls gene expression across all domains of life (1). In the context of bacterial genetic system development, reliable promoters are essential to drive genetic payload expression in the organism of interest (2,3). Constitutive promoters with constant, predictable activity are particularly desirable as they remain active across growth phases and between conditions (4,5). Yet, most promoters have not been characterised for the majority of bacterial species (6,7). This increases the number of variables that require optimisation and testing during genetic system development and remains a substantial bottleneck for the engineering of non-model microorganisms.

Bacterial transcription is regulated by sigma factors, which reversibly bind RNA polymerase (RNAP) and target it to specific promoter sequences (8). All bacteria express at least one σ^70^ family protein called the primary, housekeeping, or vegetative σ factor, which is responsible for the expression of most or all unconditionally essential genes (9). This enzyme recognises a pair of hexamer motifs (TTGACA (−35) and TATAAT (−10)) that are positioned approximately 10 and 35 nucleotides upstream of the transcription start site (TSS) and are separated by a 15 to 19 base-pair spacer (10,11). In some cases, promoter activity can also be influenced by an extended −10 (TGn) hexamer that compensates for degenerate or absent −35 sequences (12). However, these consensus features were defined in model Gammaproteobacteria. Recent work in more divergent lineages, particularly those with atypical genomic GC content, has revealed significant variability in these hexamer motifs, complicating promoter prediction outside well-studied organisms (13,14).

Over the past two decades, considerable effort has been directed toward the development of computational strategies for bacterial promoter prediction. Tools have employed position weight matrices to capture conserved motifs (15), support vector machines to classify promoter versus non-promoter sequences (16), and more recently, convolutional neural networks (CNN) to learn regulatory features directly from data (17). Despite these advances, tool performance often degrades when applied to organisms that are poorly represented in their training data (18). Moreover, even when true promoters are successfully identified, these methods provide no quantitative information about their likely activity levels.

Improving predictive models requires experimental data from diverse microbes, yet generating this data relies on functional genetic systems with reliable promoters (19). To break this bottleneck, an intermediate strategy is needed to accelerate genetic tool creation across the bacterial domain without demanding exhaustive prior characterisation.

Here we present ProFinder, a computational method that predicts a ranked shortlist of constitutively active σ^70^ promoter candidates from any bacterial genome assembly. Instead of attempting to predict all promoters in an input genome, ProFinder searches only the intergenic regions upstream of conserved, stably expressed marker genes and returns up to ten ranked promoter candidates, trading exhaustive recovery for a reliable shortlist that are ready for immediate testing in genetic constructs. We benchmark ProFinder’s precision against four existing genome-wide predictors in *Escherichia coli* and *Bacillus subtilis*, validate its predictions against published TSS-mapped promoter lists in 24 species spanning seven bacterial phyla, and apply the method to 56,742 representative bacterial genomes to generate a pan-bacterial catalogue of putative constitutive promoters for immediate use by the community.

## Materials and Methods

### HMM profile collection and screening

HMM profiles for bacterial housekeeping and phylogenetic marker genes were collected from ten public databases (339 files, 1,244 profiles; Table S1). Profiles with identical model content were collapsed to a single representative, leaving 838 unique profiles (Tables S2, S3). The 838 profiles were searched with HMMER v3 (20) against the proteomes of five reference bacteria: *Escherichia coli* K-12 MG1655 (GCF_000005845.2), *Bacillus subtilis* 168 (GCF_000009045.1), *Mycobacterium tuberculosis* H37Rv (GCF_000195955.2), *Pseudomonas aeruginosa* PAO1 (GCF_000006765.1) and *Salmonella enterica* serovar Typhimurium LT2 (GCF_000006945.2). A single permissive threshold of 25 was used for every profile to enable gene family detection across diverse bacteria; specificity was instead imposed by the filtering strategy described below.

### Marker panel definition

Expression data were taken from published compendia for the same five organisms: the *E. coli* (2,710 conditions), *B. subtilis* (265) and *M. tuberculosis* (647) modulomes, the *P. aeruginosa* PRECISE-411 dataset (411) and the *S. enterica* core compendium (533) (21). To determine expression breadth, the fraction of conditions exceeding a candidate expression threshold was tabulated across thresholds from −8 to +2 log_2_-TPM in steps of 0.25 for each organism. At each threshold we counted the families that passed in at least 95% of conditions in all five organisms simultaneously. Filtering by expression magnitude was performed by defining each family’s expression floor as the lowest of its five per-organism median values. Families were ranked by that floor and the ranking cut at its elbow, located with the Kneedle algorithm (22). Panel stability was tested by repeating both filters five times, each time withholding one organism and re-deriving the magnitude cut-off from the reduced panel, so each run applied its own threshold. The six sets (five leave-one-out runs and the full run) were compared by pairwise Jaccard similarity and by families gained and lost. Those retained by the full run but not by every leave-one-out run were discarded, leaving the families that qualify under the strictest threshold produced by any run.

### Expression dispersion and generalisation

Expression dispersion was measured for every conserved family in every organism as the difference between the 97.5th and 2.5th percentiles of centred log_2_-TPM values across conditions. Panel families were compared with rejected families by Mann–Whitney U test, pooled, within each organism, and within quartiles of expression level so that any difference could not be an artefact of expression magnitude alone. Canonical housekeeping classes (ATP synthase, RNA polymerase, ribosomal proteins, translation factors, chaperones) were assigned from HMM profile names and summarised separately. Generalisation beyond the five defining organisms was assessed in six organisms that were not used for panel selection (*Pseudomonas putida*, *Pseudomonas syringae*, *Staphylococcus aureus*, *Streptomyces coelicolor*, *Synechococcus elongatus* and *Streptococcus pneumoniae*), using published expression compendia for each (21). Panel and rejected families were located in each proteome by HMM search as above. Because the seven datasets differ in scale and normalisation, each family’s dispersion was converted to a rank within its own organism, expressed as a percentile in which 0 is the narrowest gene in that organism, so that values are comparable across datasets. Panel and rejected families were compared by two-sided Mann–Whitney U test on per-family median percentiles; the per-organism difference was tested across organisms by a two-sided Wilcoxon signed-rank test paired by organism, and each panel family’s mean percentile across organisms was tested against the median gene by a two-sided one-sample Wilcoxon signed-rank test.

### Transcription start sites and intergenic regions

TSS coordinates for 49 bacteria were taken from a published compilation (6). For each organism the reference chromosome and GFF3 annotation were retrieved under the accession given by the authors and coordinates were checked against the assembly used in that work. Intergenic regions (IGRs) were defined as the intervals between consecutive annotated genes on the same contig. For this analysis every annotated gene feature was used as a boundary, including tRNA, rRNA, ncRNA and tmRNA genes, so that an interval counted as intergenic is not in fact a transcribed structural RNA gene. IGRs were classified by the orientation of their flanking genes as tandem+ (→ IGR →), tandem− (← IGR ←), divergent (← IGR →) or convergent (→ IGR ←). The first three lie immediately 5′ of at least one start codon and are grouped here as upstream-of-start-codon (USS); convergent IGRs, which cannot carry a promoter for either flanking gene, form the complementary class and serve as a negative comparator. Intervals at contig ends were excluded.

Each TSS was assigned either to a gene body or to an IGR by coordinate overlap against the same merged gene track, so that the two classes are complementary by construction; intergenic TSSs were then assigned to individual IGRs by interval lookup. TSS density (TSSs per kb) was computed over merged gene length and over total IGR length, both pooled across the 49 genomes and separately for each genome, and fold enrichment was taken as the ratio of the two. USS and convergent IGRs were compared in the same way, as a per-genome density ratio and across genomes by a one-sided Wilcoxon signed-rank test (alternative: USS > convergent), paired by genome. Per-IGR occupancy, the fraction of IGRs carrying at least one TSS, was computed for USS IGRs in length bins of ≤ 40, 41-60, 61-80, 81-100, 101-150, 151-300, 301-500, 501-1,000, 1,000-2,000, > 2,000, each with a 95% Wilson score interval. The fraction of all TSS-containing USS IGRs falling in the 81–1,000 bp range was computed to assess that range as a selection window.

### σ^70^ motif catalogue

Per-organism σ^70^ −10 and −35 hexamers from experimentally mapped TSSs in the same 49 genomes were taken from prior work (6). Within each genome, promoters were clustered on their hexamer pair. Each pair was encoded as a one-hot vector over 12 positions, clustered by *k*-means with *k* chosen per genome from 2 to 8 by silhouette score. Subgroups from any genome sharing the same consensus (−35, −10) hexamer pair were then pooled into a single position weight matrix (PWM) by averaging their constituent matrices. Sharing across taxa was quantified per genome as the fraction of that genome’s subgroups whose consensus also occurs in at least one other genome, computed separately for the −10 hexamer, the −35 hexamer and the complete pair, and both unweighted and weighted by the number of promoters each subgroup represents. Conservation of each element was summarised as the number of distinct consensus hexamers in the catalogue and the share of promoters carried by the most common ones. Whether hexamer usage follows ancestry or base composition was tested by partial Mantel tests, run separately for the −10 and −35 elements. A between-genome distance in hexamer usage was compared with a taxonomic distance while holding difference in genome GC content constant, and with GC distance while holding taxonomy constant, with significance from matrix permutation. The −35-to-−10 spacer length was measured as a median per genome and as the share of promoters with a spacer of 16–18 bp. The distance from the −10 element to the start codon of the downstream gene was measured as a median per genome.

### Catalogue validation and choice of search window

The motif catalogue was validated by leave-one-genome-out. For each of the 49 genomes, the subgroups of the other 48 were pooled by consensus hexamer pair into a fold-specific catalogue of 65–72 motifs, which was scanned against the held-out genome’s upstream intervals using that genome’s own base composition. A candidate required a −10 at *p* ≤ 1e-2 and a −35 16–18 bp upstream of it, with the −10 within 300 bp of the downstream start codon, and was assigned to the first of three rules it met: path A, the −35 catalogued with that −10 at *p* ≤ 1e-3; path B, the same pairing with the −35 relaxed to *p* ≤ 3e-2 where the −10 carried an upstream TG dinucleotide (the extended −10, which compensates for a weak −35 (23)); or path C, the best −35 from any other catalogued pair at *p* ≤ 1e-2, Šidák-corrected for the number of pairs searched (24). The three rules therefore differ only in which −35 they accept. These p-value thresholds were selected as the tightest setting for the majority of genomes to return at least ten promoter candidates without a loss of accuracy. Promoter recovery was measured on intervals carrying an experimentally mapped TSS as the fraction with a complete −10/−35 pair under at least one rule, and compared with either upstream intervals carrying no mapped TSS, or dinucleotide-matched null sequence. Both comparisons are reported pooled across folds and per genome.

### The ProFinder pipeline

Coding sequences are first called with Prokka (25). Intervals between consecutive coding sequences are kept at 80–1,000 bp, and only tandem+ and tandem− intervals are carried forward. Operons are grouped from same-strand coding sequences separated by no more than 25 bp and bounded by intervals of at least 75 bp, and are kept only when exactly one end is gene-proximal, so that one upstream interval can be assigned unambiguously. An operon is marker-associated if any of its coding sequences hits one of the 98 bundled profiles at a full-sequence bit score of at least 25. A candidate promoter requires a −10 hit whose 3′ edge lies within 300 bp of the downstream start codon and a −35 hit at a spacer of 16–18 bp, and is assigned to the first matching of paths A, B and C as defined above. Each interval is scored with the 72 paired PWMs as log-odds against the A/C/G/T frequencies of the input assembly itself, so that no genome is scored against another genome’s composition. Motif significance is assessed against a null of dinucleotide-preserving shuffles of intervals matched for length and composition and the thresholds applied to the −10 and −35 *p*-values. Candidates are ranked by a logistic model trained on the 49-genome TSS panel to predict whether an interval contains an annotated intergenic TSS, with intragenic sense and antisense starts not counted as positives; its 14 inputs summarise motif significance and rank, classification path, interval and spacer geometry, and base composition.

### Benchmarking in model organisms

ProFinder was compared with PromoterLCNN (26), Sigma70Pred (27), SAPPHIRE.CNN (28) and Promotech (29), each under its default settings. Two ProFinder outputs were used: the full motif-confirmed set of tandem IGRs, with no marker-gene filter and no shortlist cut-off, and the ranked shortlist of ten, obtained by keeping intervals that are both marker-associated and motif-confirmed and taking the ten with the highest rank score. PromoterLCNN was given the IGR set extracted by ProFinder for the same organism and its σ^70^-classed calls taken as its output. Sigma70Pred, SAPPHIRE.CNN and Promotech are genome scanners and were run on the whole assembly. Calls were validated against the experimentally characterised promoters of *E. coli* K-12 MG1655 (RegulonDB (30)) and the union of the DBTBS (31) and SubtiWiki (32) records for *B. subtilis* 168 using BLASTN with a default both-strand search. Precision is reported with 95% Wilson score intervals, and tools were compared within each organism by Fisher’s exact test across all pairs with Benjamini–Hochberg correction. Because PromoterLCNN does not rank its calls, its output was filtered to ^70^ calls upstream of marker genes and resampled as random draws of ten, from which we took the mean number of false positives per ten and its 95% range.

### Cross-species validation

ProFinder and PromoterLCNN were run on 24 genomes (Table S33) bacterial genomes chosen to span the bacterial phyla and a GC range of 29–72%. Both tools received the identical tandem IGR set for each genome. For each genome we recorded the number of tandem IGRs, ProFinder’s marker-associated motif-confirmed candidates and its shortlist of ten, and PromoterLCNN’s σ^70^ calls. Per-IGR yield was defined as calls divided by tandem IGRs, summarised by mean, standard deviation, coefficient of variation, range and maximum-to-minimum fold span, and related to genome GC content by Pearson *r* and Spearman ρ for each tool separately. Shortlisted promoters were then scored against each organism’s published TSS-mapped promoter list (Table S33). A shortlisted interval counted as validated when a reference TSS fell inside the predicted interval. The rate expected without signal was estimated per genome from composition-matched intervals, and each genome’s observed rate was tested against its own null by a binomial test. ProFinder’s shortlist of ten was compared with random draws of ten from PromoterLCNN’s ^70^ marker-gene calls for the same genome, using the same resampling as above, and the paired difference across genomes was tested by a Wilcoxon signed-rank test.

### Expression validation at extremes of genome GC content

The two highest-and two lowest-GC genomes of the cross-species panel (*Streptomyces coelicolor* A3 (2), *Caulobacter crescentus* NA1000, *Campylobacter jejuni* NCTC11168 and *Clostridioides difficile* 630) were validated against their own transcriptomes. Public Illumina RNA-seq runs for these four taxa were listed from the European Nucleotide Archive (3,891 runs) and filtered to runs of at least 10□ reads whose metadata matched the reference strain, excluding library types that do not measure transcript abundance. Sample titles were normalised to collapse replicates, one run was kept per distinct condition, at most six runs were taken from any one study, and genetic perturbations were capped at two per organism, giving 32 runs from 18 studies. The first 2 × 10^6^ reads of each run were quantified with Salmon v2.7.0 (33) against coding sequences extracted from each genome’s GenBank annotation. Runs were discarded if fewer than 30% of coding sequences were detected or if fewer than 50,000 reads were assigned, leaving 29 conditions, 6–8 per organism.

A gene was counted as expressed if it exceeded 1 TPM in at least half the retained runs of its organism, and as expressed in every condition if it exceeded 1 TPM in all of them. Its dynamic range was taken as log_2_ of the ratio of the 97.5th to the 2.5th percentile of its TPM across conditions. Panel members were located by searching the 98 bundled profiles against each proteome at a bit score of at least 25 and keeping the top-scoring protein per family. Panel genes were compared with all other expressed genes of the same organism, for dynamic range and for median expression, by Mann– Whitney test. Genes immediately downstream of shortlisted promoters were compared with 2,000 size-matched random draws from the expressed set. Finally, the panel’s dynamic range in the two high-GC organisms was compared with that in the two low-GC organisms, and yield against GC content across the four organisms by Spearman ρ.

### Domain-wide application

ProFinder was applied to 56,742 bacterial species drawn from a MAG collection of animal-, human-and environmental-associated assemblies (34). Representatives were chosen on the collection’s own quality score. Each genome was run with the default ProFinder parameters. Per-genome output tables were concatenated and joined to GTDB taxonomy from phylum to species, genome GC content and genome size. Genomes returning a shortlist of 10 promoters were classed as complete and the remainder incomplete, and genome-size and GC distributions were summarised separately for the two groups. For each phylum with at least 20 genomes, the incomplete fraction was compared with the domain-wide rate by a binomial test with Benjamini–Hochberg correction across phyla. Shortlist completeness was related to GC content and to genome size by Spearman correlation, and medians were also computed within evenly spaced GC and genome-size bins to detect non-monotonic trends. Spacers within the 16-18 bp window were pooled and for each phylum contributing at least 100 calls, tested against equal use of the three permitted spacings by a chi-squared goodness-of-fit test. The genes carrying the calls were ranked by how often each gene symbol was served, and concentration reported as the number of symbols accounting for half and for 80% of all calls. These are annotation symbols on the served genes, not members of the 98-profile marker panel; the operon filter admits every gene of a qualifying operon, so a served gene need not itself be a panel family.

### Statistical analysis

Tests are two-sided unless stated otherwise. Proportions are reported with 95% Wilson score intervals. Where a family of related comparisons was made, *p-*values were adjusted by the Benjamini–Hochberg procedure, and the family over which the adjustment was applied is stated with each result. Analyses used Python 3 with NumPy, SciPy and pandas; HMM searches used HMMER v3 and sequence comparisons NCBI BLAST+.

## Results

### A conserved marker gene panel enriched for stable expression across bacterial phyla

To identify a panel of marker genes that are both broadly conserved and expressed with limited dependence on growth condition, we screened HMM profiles of known marker genes (*n* = 838; Tables S1-S4) against five reference bacterial species spanning Gammaproteobacteria, Bacillota, and Actinomycetota (*E. coli*, *B. subtilis*, *M. tuberculosis*, *P. aeruginosa*, and *S. enterica*) (Fig. 1A; Table S4, S5). These organisms were selected because large, condition-rich transcriptomic compendia were available for each. This screen identified 261 gene families that were conserved in every organism (Table S5).

**Figure 1.**
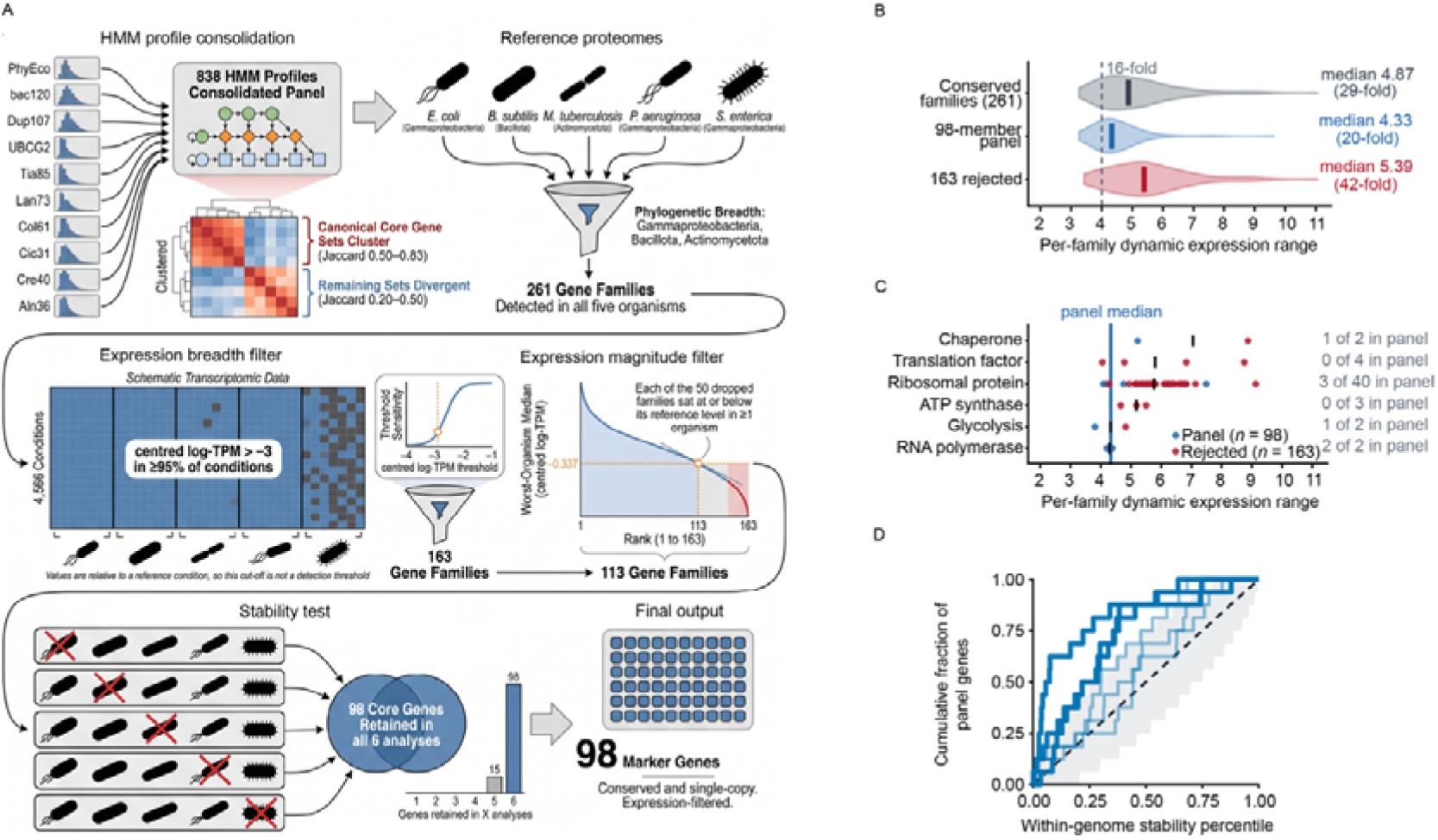
Construction of the organism-independent marker panel and validation of its expression behaviour. (A) Schematic of panel construction; the number of gene families surviving each stage is given in the diagram. (B) Distribution of per-family dynamic expression range for the conserved families entering the expression filters, the families retained in the panel and those rejected. Dynamic range is the log2 span between a family’s 2.5th and 97.5th expression percentiles across conditions, evaluated in whichever of the five organisms it is widest. Violins show the full distribution, coloured bars the group medians, annotated with the equivalent fold-change, and the dashed line a 16-fold reference span. (C) The same statistic for families assigned to six canonical housekeeping classes, one point per family, coloured by whether the family was retained in the panel or rejected. Black bars mark class medians, the vertical line marks the median of the retained panel, and right-hand labels give the number of families of each class retained out of the total in that class. (D) Expression stability of the panel in six organisms not used to define it. Cumulative distribution of the within-genome stability percentile of the panel families present in all six organisms. The percentile ranks each gene among all genes of its own genome by the residual of its expression standard deviation after the genome-wide mean–variance trend is removed, so 0 is the most stable gene in that genome and 0.5 the median gene. One step curve per organism, drawn solid where that organism’s genes alone depart from the uniform expectation at a two-sided Kolmogorov–Smirnov p < 0.05 and faint where not. Dashed diagonal, the expectation for genes drawn at random from the same genome; shaded band, the pointwise 2.5th–97.5th percentile interval of that expectation from 20,000 simulated draws of 16 genes.

We then sought to identify the gene families that were consistently expressed in all organisms across different experimental conditions. For this, we mapped gene expression from 4,566 conditions including diverse cultivation media, stresses, antibiotic exposures, genetic perturbations, and growth phases to determine a minimum expression threshold (Fig. 1A; Table S6) (21). The number of genes retained decreased as the log_2_-TPM threshold was increased from −4 (228 families) to −1 (none), with a discernible elbow at −3 which we adopted as the minimum threshold (Table S7). We then ranked the 163 surviving families by their lowest per-organism median and cut at the inflection point (−0.34, a 1.3-fold reduction), which left 113 families (Fig. 1A). These represented the 113 gene families that were actively expressed in most conditions across all organisms tested.

To ensure that the 113 families were stable across organisms we repeated the selection five times, leaving one bacterium out each round and keeping the cut-offs unchanged. All 113 survived every round, while the number of additional qualifying families changed. For example, there were 81 more without *E. coli*, whose 2,710 conditions made it the hardest test to pass, compared to 16 without *B. subtilis*, six without *S. enterica* and none without *M. tuberculosis* or *P. aeruginosa* (Fig. 1A; Tables S8, S9). Without *E. coli*, the recalculated cut-off was stricter (*n* = 98) and we adopted this number as the final, more conservative panel definition. These 98 genes represented those that remained active in almost every condition tested, in all five bacteria at once, making their promoters promising candidates that work regardless of growth condition (Table S10).

To determine each family’s expression variance across the dataset, we computed the span between the 2.5th and 97.5th percentiles of its centred log_2_ expression in each organism. Across the 261 conserved families, expression spanned a median of 29-fold, with the 98 panel families representing the narrow tail of that distribution at 20-fold, compared to 42-fold for the families the filters rejected (two-sided Mann–Whitney *p* = 1.5e-12; Fig. 1B, Table S11). Several canonical housekeeping classes showed the widest expression range overall: ribosomal proteins spanned 54-fold with 3 of 40 families retained, translation factors 56-fold with none of four, and chaperones 132.1-fold with one of two (Fig. 1C), indicating that housekeeping annotation is not on its own a reliable predictor of constant expression. Therefore, the 98-gene panel represents the low-variance tail of a conserved gene repertoire.

To test whether the panel behaves the same way in bacteria that played no part in selecting it, we examined six further organisms (*P. putida*, *P. syringae*, *S. aureus*, *S. coelicolor*, *S. elongatus* and *S. pneumoniae*), ranking every gene in each by how variable its expression was. Panel genes ranked as more stable than the equally conserved families the filters had rejected (median percentile 0.30 versus 0.40; Mann–Whitney *p* = 0.0056), in all six organisms (Tables S12, S13), and every panel family was more stable than its genome’s median gene (Fig. 1D). Their expression varied about 15% less than expected for genes of their abundance in every organism, and by a third in *S. coelicolor*. Therefore, the panel genes chosen in five bacteria behaved the same way in six others. Together, these results define a 98-gene panel that can be identified from genome sequence alone whose transcription is maintained across conditions wherever it is found, providing a robust anchor set for predicting their promoters.

### Transcription start sites concentrate in intergenic regions upstream of start codons

We then determined whether non-coding intervals between genes (intergenic regions (IGRs)) were enriched for promoters across diverse bacteria. Using 67,005 experimentally characterised transcription start sites (TSSs) from 49 bacterial genomes (Table S14, S15) (6), we found that IGRs contained 76.8% of all TSSs while constituting only 12.0% of total chromosomal DNA (51,484/67,005; Fig. 2A; Table S16). This corresponded to a 24-fold density enrichment over coding sequence (2.26 vs. 0.093 TSSs/kb), and this enrichment was consistent in every genome (median: 56-fold; interquartile range 22–93-fold). TSS location was additionally dependent on IGR type. IGRs positioned upstream of a start codon (USS), comprising tandem+, tandem− and divergent IGRs, carried higher TSS densities than convergent IGRs in all genomes (Fig. 2B, C). Together, this supported the expectation that promoters are predominantly located in IGRs that are located upstream of genes.

**Figure 2.**
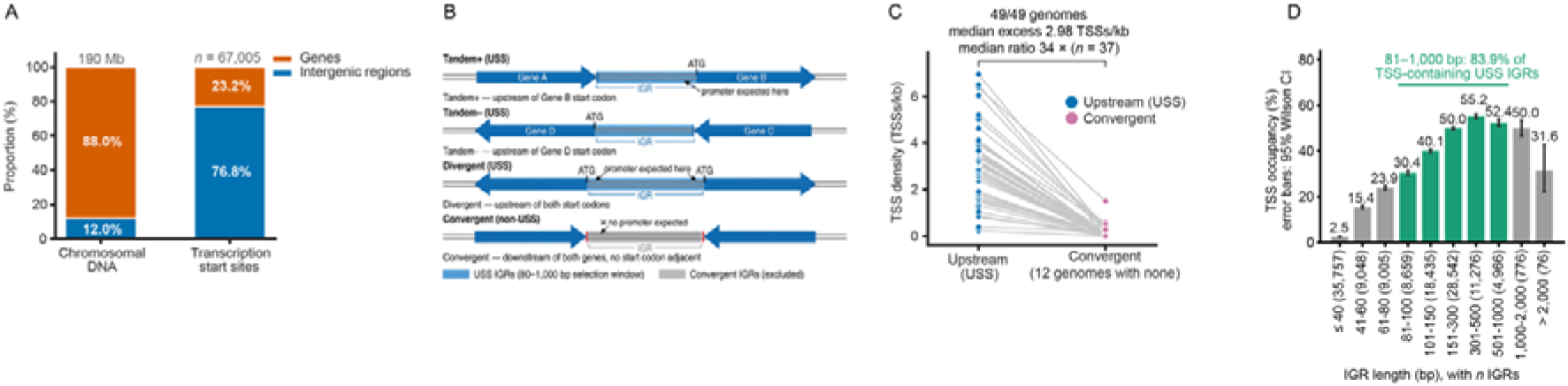
Transcription start sites concentrate in a definable subset of intergenic regions. (A) Composition of the 49 panel genomes, pooled. Left, the proportion of total chromosomal DNA that is intergenic; right, the proportion of mapped transcription start sites that fall in intergenic sequence. Intergenic regions are the gaps between consecutive annotated gene features. (B) Schematic of the four intergenic-region classes defined by the orientations of the two flanking genes. Tandem+ and tandem− regions lie immediately upstream of one flanking start codon and divergent regions upstream of both, so a promoter is expected in each; these three form the upstream-sense set (USS) used throughout this figure, within a length window of 81–1,000 bp. Convergent regions lie downstream of both flanking genes with no adjacent start codon and are excluded. (C) Transcription start site density, as sites per kb of intergenic sequence, in upstream-sense against convergent regions. One point per genome in each class, joined by a line so the comparison is read within a genome. Annotations give the number of genomes in which upstream-sense density is the higher of the two, the median paired difference, and the median of the per-genome ratio. (D) Percentage of upstream-sense intergenic regions containing at least one mapped start site, by region length. Bar colour distinguishes bins lying inside the 81–1,000 bp selection window from those outside it, the number of regions per bin is given on the axis, the value above each bar is that bin’s occupancy, and error bars are 95% Wilson confidence intervals. The bracket gives the share of all start-site-containing upstream-sense regions that fall within the selection window.

The proportion of USS IGRs containing a detected TSS increased monotonically with interval length: 2.5% at ≤ 40 bp, 15.4% at 41–60 bp, 23.9% at 61–80 bp, 30.4% at 81–100 bp and 40.1% at 101–150 bp (Fig. 2D). Occupancy peaked at 55.2% in the 301–500 bp bin and then decreased (52.4% at 501– 1,000 bp; 50.0% at 1,001–2,000 bp), and the 81–1,000 bp window captured 83.9% of all TSS-containing USS IGRs (19,233/22,927; Fig. 2D). Only 1.1% of TSS-bearing USS IGRs exceeded 1,000 bp. Beyond 2,000 bp 76 USS IGRs remained, which were too few to estimate occupancy precisely (31.6%, 95% CI 22.2 -- 42.7%). Thus, USS IGRs combined with a length range < 1,000 bp provided a conserved geometric criterion for detecting TSSs in every genome examined.

### A cross-species catalogue of ^70^ promoter architecture

We next asked how much of ^70^ promoter architecture was shared across bacteria. Clustering the −10 / −35 hexamer pairs already mapped in the 49 genomes (6) gave 72 distinct pairings, or architectures, covering 66,999 promoters (Tables S17, S18). Few were widely used, with the nine most common architectures accounting for 51.8% of all promoters and the 26 most common for 80.7% (Fig. 3A). The common architectures were also the widespread ones, for example only 17 of the 72 appeared in more than one genome, yet those 17 included eight of the nine most common (Fisher’s exact *p* = 1.6e-5; Fig. 3B). Each genome was similarly concentrated, with a single architecture accounting for a median 52.6% of its promoters and for at least half in 38 of the 49 (Table S19). In 35 genomes that most-used architecture was shared with at least one other genome, while in the remaining 14, including *B. subtilis*, *M. tuberculosis*, *P. aeruginosa* and *S. coelicolor*, it was found nowhere else. A few dozen architectures therefore account for most promoters in most genomes, though a substantial minority of genomes are dominated by architectures unique to them.

**Figure 3.**
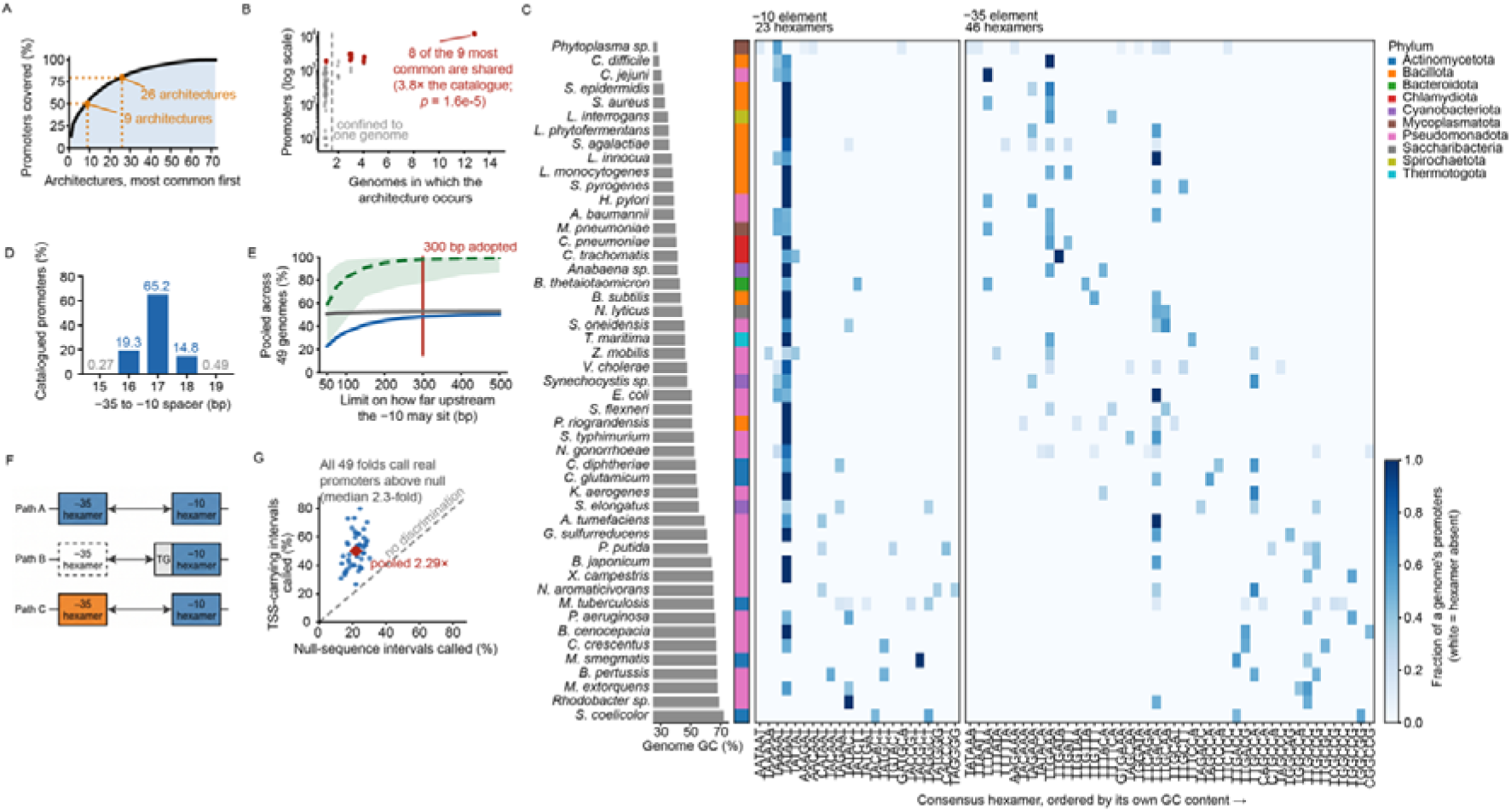
A shared σ^70^ motif catalogue. (A) Cumulative share of catalogued promoters accounted for by architectures ranked from most to least abundant, an architecture being a distinct pair of consensus −35 and −10 hexamers. Guides mark the number of architectures reaching half and 80% of promoters. (B) For each architecture, the number of promoters it carries against the number of genomes in which it occurs, on a logarithmic count axis. Architectures confined to a single genome are separated by the dashed line and shown in grey; the most abundant architectures are highlighted. The annotation gives how many of the most abundant are found in more than one genome, the enrichment of shared architectures among them relative to the catalogue as a whole, and a Fisher exact test of that enrichment. (C) Per-genome use of each consensus hexamer. Rows are the 49 catalogue genomes ordered by genome GC content, with the adjacent bar giving that value and the colour strip the phylum. The two heatmaps show the 23 distinct −10 and 46 distinct −35 consensus hexamers; columns are ordered by the hexamer’s own GC content. Cell colour is the fraction of that genome’s promoters carrying the hexamer, and white indicates the hexamer is absent from that genome. (D) Distribution of the −35 / −10 spacer over all catalogued promoters of the 49-genome panel. Bars give the percentage of promoters at each spacer length observed, labelled above; lengths of 16–18 bp are shaded and those outside that range are grey. (E) Effect of the limit on how far upstream of a start codon the −10 element may sit, swept from 50 to 500 bp and evaluated on the same 49 genomes. Three quantities are plotted against that limit, all as percentages: Reachable (dashed line) is the percentage of TSS-bearing intergenic intervals holding at least one catalogued promoter whose −10 falls within the limit. The shaded band spans the lowest and highest values among the 49 genomes. Recovered (solid blue) is the percentage of the same intervals in which the scan placed a complete −10 / −35 pair, scored leave-one-genome-out. Correct (solid grey) is the percentage of intervals called at that limit that are TSS-bearing. The vertical line marks the limit adopted throughout this work. Lines are pooled over the 49 genomes. (F) Schematic of the three gating paths. Every path requires a −10 and a −35 element separated by the permitted spacer and differs only in which −35 is accepted. The dashed outline marks the relaxed threshold. Paths are tried in order and a candidate is assigned to the first it matches. (G) Held-out transfer of the motif catalogue. Each point is one leave-one-genome-out fold: the percentage of that genome’s TSS-carrying intergenic intervals in which the gate placed a complete −10 / −35 pair, against the percentage of dinucleotide-matched null intervals called in the same genome. Dashed diagonal, equal call rates. Red diamond, the pooled rate across all folds.

The two halves of these architectures were not equal, with the –10 motif more conserved than the –35 element. Overall, there were only 23 different −10 sequences compared to 46 different −35s (Fig. 3C; Table S20, S21). The canonical –10 motif, TATAAT, dominated and was situated upstream of 67.1% of all promoters, while the most common −35 motif, TTGACA, covered only 19.4% (Table S22). Specifically, TATAAT was combined with 28 different −35 sequences, while other −10 motifs were often associated with a single –35 partner (14 of 23). This hexamer distribution was not phylogenetically structured (partial Mantel *r* = −0.04 and +0.02, *p* ≥ 0.59), but was explained by genome composition independently of phylum (*r* = +0.27 and +0.26, *p* ≤ 0.0003), with –35 motif GC content scaling with genome-wide GC content (Fig. 3C; Table S23). Thus, σ^70^ architecture is largely asymmetric with the −10 element more conserved than the −35 across the bacterial tree, and where there is variation it tracks genome composition rather than descent.

The spacer regio separating the −35 / −10 sites was near-invariant, with a median of 17 bp in every genome tested and in 99.2% of promoters it fell within 16–18 bp (Fig. 3D; Table S24). The distance from the −10 element to its gene was less conserved with a median of 63 bp across the catalogue, while per-genome medians ranged from 35 to 204 bp (Table S25). To determine whether one window could accommodate that variation, we swept the limit on this distance from 50 to 500 bp, scanning each genome’s intergenic intervals with a catalogue rebuilt without it. At 100 bp, the proportion of TSS-bearing intervals holding a promoter spanned 46% to 97% between genomes, but by 300 bp that proportion was 98.0% overall and above 77% in every genome (Fig. 3E; Table S26, S27). Within 300 bp, the fraction of intervals containing a −10 / −35 pair increased from 22.9% to 48.4%, at no cost in precision (50.8% to 52.9%; Fig. 3E). Thus the −10 / −35 spacer is essentially fixed across the bacteria tested while the promoter’s distance from its gene is not, and a 300 bp window accommodates that variation.

We then asked whether the catalogue could detect −10 / −35 hexamers in genomes that had not contributed to it. Given the variability of the −35 element, we combined three alternative −35 acceptance rules into a single gate (see Materials and Methods; Fig. 3F). To test it we rebuilt the catalogue 49 times, each time excluding one genome before clustering, so that neither the architecture definitions nor the PWMs had seen the genome being scanned, and calibrated each scan against that genome’s own base composition. The gate additionally required 16–18 bp between the −10 and −35 hexamers and no more than 300 bp between the −10 element and the start codon. Applied to each held-out genome, complete hexamer pairs were placed in 50.1% of TSS-bearing intergenic intervals, a 2.05-fold enrichment over intervals with no mapped TSS and 2.29-fold over dinucleotide-matched null sequence (Fig. 3G). Both comparisons held in all 49 genomes individually (per-genome median 2.04-fold against TSS-negative intervals, range 1.33–4.30, and 2.30-fold against null, range 1.22–3.97; paired Wilcoxon *p* = 3.6e-15 for each). Whether the catalogue had seen a genome made almost no difference (50.1% compared to 50.6%), suggesting that −10 / −35 motifs can be identified in genomes with uncharacterized promoters.

### Constitutive promoter prediction with ProFinder

We integrated these layers into an end-to-end computational method for candidate constitutive promoter prediction in bacteria: (i) a 98-gene marker panel for predicting conserved, putative constitutively expressed genes; (ii) coding sequence (CDS)-adjacent USS IGR geometry; and (iii) the 72-PWM σ^70^ motif catalogue with spacing constraints. Named ProFinder, this pipeline represents an end-to-end strategy that takes a raw bacterial genome and returns a ranked shortlist (up to 10) of putative constitutive promoters for use in genetic tool development for diverse bacteria (Fig. 4A).

**Figure 4.**
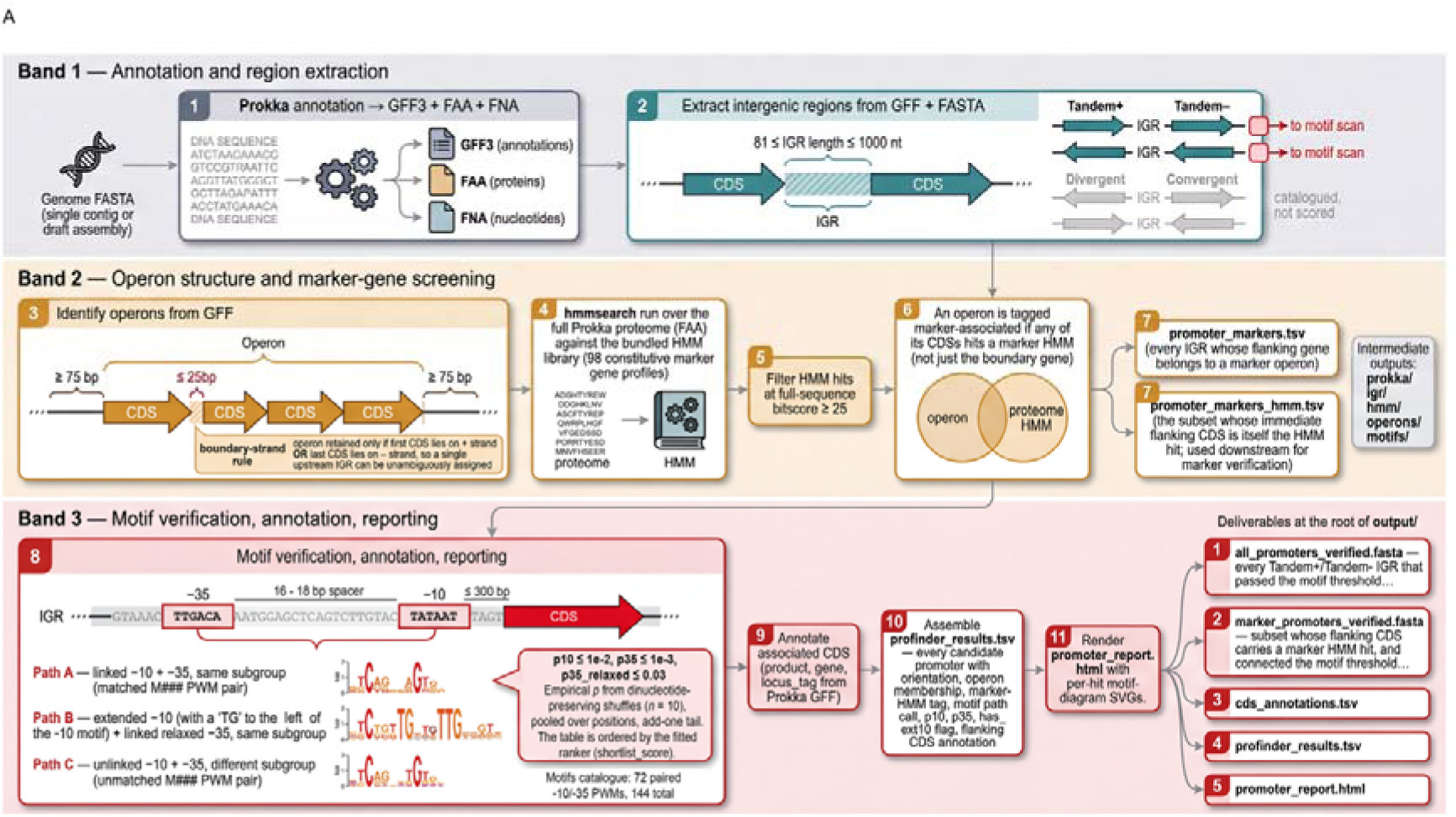
The ProFinder pipeline. (A) The computational pipeline carried out by ProFinder, from an input genome assembly to a shortlist of candidate constitutive promoters.

First, coding sequences are predicted and annotated with Prokka (25) and intergenic regions in the 81−1000 bp range are extracted and classified by flanking gene orientation. Only tandem+ and tandem-IGRs are retained as they are more likely to contain single, one-directional promoters that are more suitable for genetic tool construction. Operon structure is predicted by grouped consecutive CDSs on the same contig when they are separated by ≤ 25 bp and flanked on either side by an IGR that is ≥ 75 bp. Operons are retained if either the first CDS lies on the + strand or the last CDS lies on the - strand (but not both), so that a single upstream IGR can be reliably assigned to each operon (Fig. 4A).

All CDSs are screened against the bundled HMM library (*n* = 98). An operon is flagged as marker-associated if at least one of its CDSs hits an HMM profile with full-sequence bit score ≥ 25, and the upstream IGR of that operon is extracted. Candidate IGRs are scanned on the 5’→3’ strand relative to the downstream gene with the 72 paired −10 / −35 PWMs from the bundled motif catalogue, scored as log-odds against the per-genome A/C/G/T background derived from the input FASTA. Each interval is compared against dinucleotide-preserving shuffles of intervals in its own length × composition stratum. Each motif hit is classified into the three paths (A, B, and C) that we defined for hexamer detection, and each detected hexamer must be separated by 16-18 bp while the right edge of every −10 hit is required to sit within 300 bp of the downstream CDS start.

Up to ten putative promoters are returned, ranked by a 14-feature logistic model that weighs motif significance, path, geometry and interval composition. ProFinder produces an output HTML report containing the candidate promoters that includes genomic coordinates, strand orientation, classification path, motif positions and their *p*-values, spacer length, ranking score, operon membership, and the associated CDS annotation. Putative promoter fasta files are also produced, in addition to output tables containing all HMM hits, IGRs, operons, and motif hits.

### ProFinder is competitive for model organisms and is predictive across phyla

ProFinder was benchmarked against four existing promoter predictors (PromoterLCNN (26), Sigma70Pred (27), SAPPHIRE.CNN (28), and Promotech (29); Table 1). A 1:1 comparison was not possible as ProFinder was specifically designed to produce a short, marker-anchored shortlist of putative constitutive promoters for specific use-cases while traditional promoter predictors are sequence or whole-genome classifiers. Therefore, we prioritised the benchmarking of positive predictive value (precision) over recall. The outputs from each tool were validated against experimentally determined promoter sets in *E. coli* and *B. subtilis* (Table S28) (30–32).

**Table 1.**
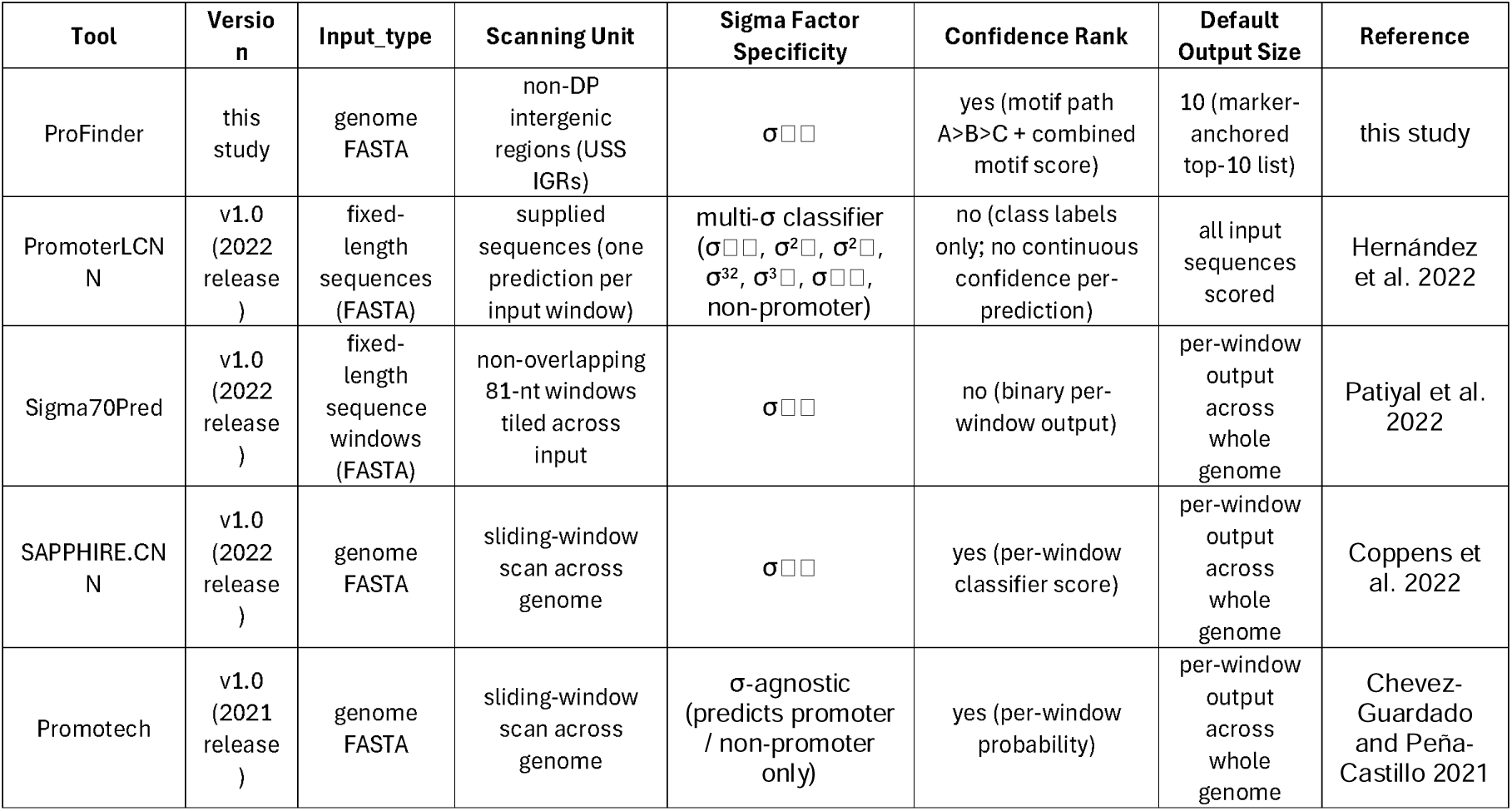
Promoter prediction tools compared in this benchmark.

Using all USS IGRs (i.e. tandem+/tandem-IGRs enriched for −10 / −35 motifs without a shortlist cut-off or marker gene filtering), ProFinder achieved 0.851 precision for *E. coli (n* = 958; false positives (FP) = 143) and 0.852 for *B. subtilis* (*n* = 1,028; FP = 152; Fig. 5A). In contrast, PromoterLCNN achieved lower precision from fewer hits for both *E. coli* (*n* = 636; precision: 0.844; FP = 99) and *B. subtilis* (*n* = 884; precision: 0.846; FP = 136; Table S29), but these outcomes were statistically indistinguishable (Fisher’s exact with BH correction, *p* = 0.749 in *E. coli* and 0.749 in *B. subtilis*). This indicated that ProFinder’s filtering criteria, geometric priors, and motif scanning were as predictive of a genuine promoter as the CNN deployed in PromoterLCNN, at least for *E. coli* and *B. subtilis.* By contrast, the remaining tools delivered larger raw outputs at lower precision (Fig. 5A).

**Figure 5.**
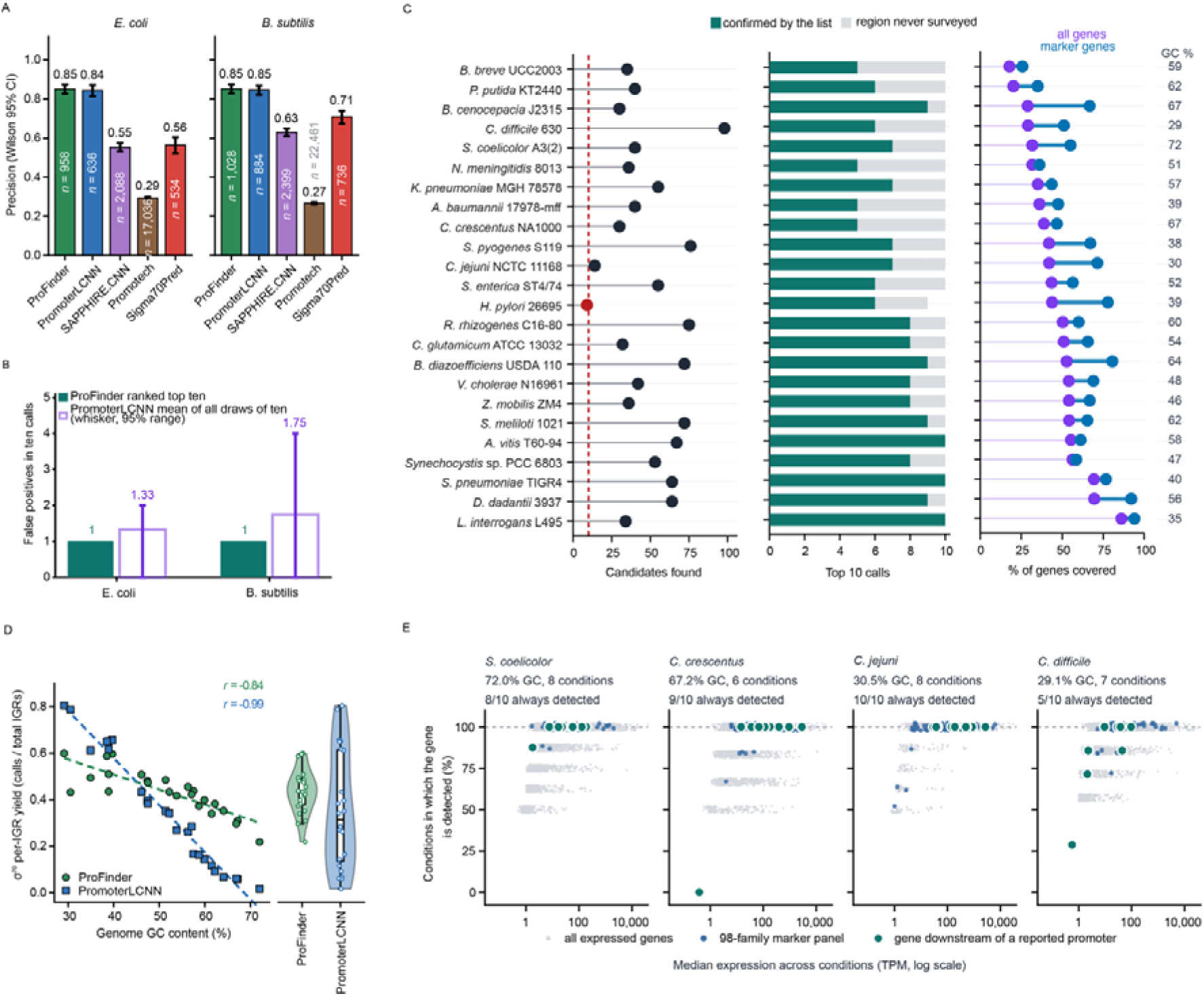
Benchmarking ProFinder against published promoter predictors and against each organism’s own promoter list. (A) Precision of five promoter predictors in E. coli and B. subtilis, scored against each organism’s published promoter list. Bar height is the proportion of a tool’s calls that are confirmed, labelled above; the number of calls scored is given inside each bar, and error bars are Wilson 95% confidence intervals. (B) False positives among ten calls. Open bars give the mean over all draws of ten and whiskers the 95% range. (C) Per-genome benchmark across 24 genomes, one row each, with genome GC content at right. From left: the number of candidate promoters ProFinder found, with a dashed line at the ten needed to fill a shortlist and the single genome falling below it highlighted; the composition of each genome’s ten highest-ranked calls, split into those confirmed by the organism’s published list and those falling in regions that list never surveyed; the percentage of genes for which the published list provides coverage, shown separately for all genes and for marker-panel genes. (D) σ70 per-intergenic-region yield against genome GC content for both tools across the same 24 genomes, with least-squares fits and Pearson correlations. Right, the distribution of each tool’s per-genome yields as a violin with interquartile box and median. (E) Expression behaviour in the two highest-and two lowest-GC genomes of the panel, one sub-panel per organism headed with its GC content, the number of RNA-seq conditions surveyed, and how many of its ten shortlisted promoters lie upstream of a gene detected in every condition. Each point is one gene, plotted as its median expression across conditions against the percentage of conditions in which it is detected, for all expressed genes, for the 98-family marker panel, and for the genes immediately downstream of a reported promoter.

When considering ProFinder’s ranked marker gene output for each organism (≤ 10 promoter candidates), a single false positive was detected for both *E. coli* and *B. subtilis*, representing a precision value of 0.90 in each case (Fig. 5B; Table S30, S31). As PromoterLCNN does not rank its predictions, size-matched random draws of ten were taken from its output after filtering to retain only putative σ^70^-regulated genes. This resulted in 1.33 false positives per ten in *E. coli* (95% range 0–2; 2 of 15 calls false) and 1.75 in *B. subtilis* (0–4; 7 of 40) (Fig. 5B). Thus, ProFinder’s ranked top 10 list produced a more accurate shortlist of putative promoters compared to a size-matched draw from the best-performing sequence-only classifier.

A further separation between tools emerged when screening a larger number of diverse organisms. Across 24 bacterial genomes spanning 29–72% GC (Table S32, S33), both tools were run on identical tandem IGR sets and scored against each organism’s published TSS-mapped promoter list. ProFinder returned a median of 41 candidate promoters per genome (range 9–98). Twenty-three of the 24 genomes yielded a full shortlist of ten (*H. pylori* yielded nine), giving 239 shortlisted promoters in total (Fig. 5C). Of these, 178 (74.5%) matched a promoter in the organism’s published TSS-mapped list (Fig. 5C). Notably, most of the unmatched fraction likely reflected incomplete reference lists rather than incorrect calls as TSS data covered a median of 43.6% of genes per organism (Fig. 5C), and 52 of the 61 unmatched calls lay in intervals with no mapped TSS at all. When an interval did carry a mapped TSS, 178 of 187 calls were confirmed (95.2%; per-genome median 100%, range 78–100%) so 74.5% precision is a lower bound. In comparison with PromoterLCNN, ProFinder validated a median 7.5 of ten compared to PromoterLCNN’s 6.25 expected across the dataset (Table S34).

Across the same 24 genomes, PromoterLCNN’s per-IGR yield varied 46.8-fold overall (range 0.0172– 0.8043, SD 0.249, CoV 0.71) compared to 2.7-fold for ProFinder (mean 0.43, SD 0.094, CoV 0.22; range 0.218–0.599; Fig. 5D; Table S35). PromoterLCNN’s variation was almost entirely determined by genome GC content (Pearson *r* = −0.99, Spearman ρ = −0.98), with its promoter yield collapsing to 0.017 calls per IGR in *S. coelicolor* (72% GC). ProFinder’s yield also declined with GC, though less steeply (*r* = −0.84, ρ = −0.86, *p* = 3e-7) and was still able to provide usable results across the full range (Fig. 5D).

To directly test whether ProFinder could identify sufficient putative constitutively expressed genes at compositional extremes, we examined the two highest-and two lowest-GC genomes of the panel: *Streptomyces coelicolor* A3 (2) (72.0% GC), *Caulobacter crescentus* NA1000 (67.2%), *Campylobacter jejuni* NCTC11168 (30.5%) and *Clostridioides difficile* 630 (29.1%) using public RNA-Seq data spanning 29 growth conditions (6–8 per organism; Table S36). Between 82 and 91 of the 98 marker families were detected per organism, 78 to 88 of these were expressed in every condition surveyed, and they together spanned a narrower expression range than other expressed genes in three of the four organisms (Fig. S1A, Table S37). They also achieved higher expression than the background in three of the four (Fig. 5E). Of the 40 shortlisted promoters, 32 were upstream of a gene expressed in every condition (Fig. 5E; Table S38), and these genes were more highly expressed than size-matched random draws from expressed genes in all four, significantly so in *C. crescentus* (145 versus 26 TPM, *p* = 0.003) and *S. coelicolor* (36 versus 9 TPM, *p* = 0.018; Fig. S1B). Thus, neither compositional extreme prevented ProFinder from filling a shortlist, and most candidates were upstream of genes expressed in every condition.

### A pan-bacterial catalogue of predicted marker gene-associated promoters

ProFinder was applied across the bacterial domain to 56,742 unique species spanning 149 phyla (Fig. 6A; Table S39, S40), GC contents of 21.4–77.0% and genome sizes of 0.28–16.04 Mb (central 98%: 27.0–73.1% and 0.67–9.35 Mb) (Fig. 6A). At least one promoter was shortlisted for 99.76% of these genomes (*n* = 56,605; 137 returned none; Fig. 6B), and a complete ten-promoter shortlist for 81.40% (*n* = 46,190) (Fig. 6A; Table S41). Coverage extended beyond model lineages, with 25 of the 28 single-genome phyla receiving a complete ten-promoter shortlist alongside well-sampled groups such as Pseudomonadota (143,338 calls) and Bacteroidota (72,122), and there was no penalty for sparsely sampled lineages (Table S42). The phyla most often producing incomplete lists were Aquificota (86.0% partial, *n* = 57), Eremiobacterota (75.5%, *n* = 155) and Margulisbacteria (68.4%, *n* = 38) (Table S43).

**Figure 6.**
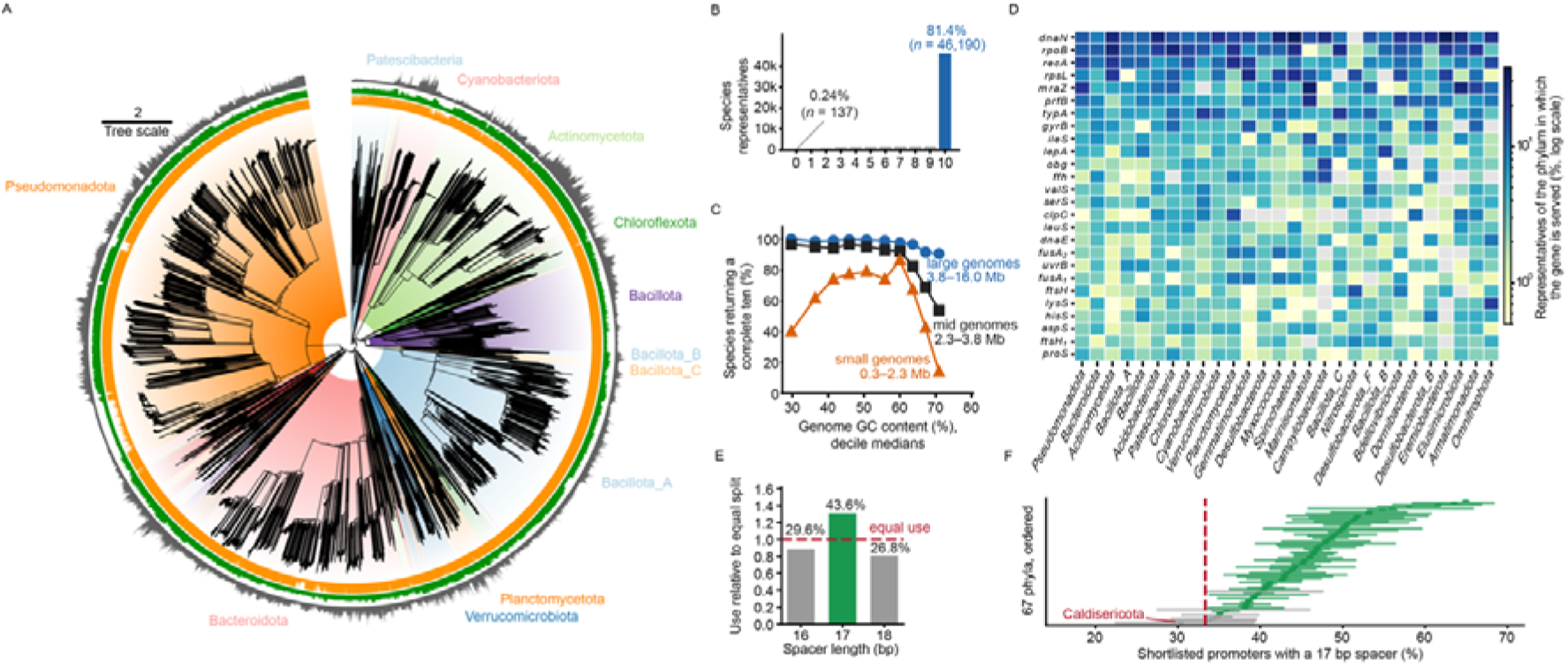
ProFinder applied across the bacterial domain. (A) Maximum-likelihood phylogeny of the bacterial species representatives analysed, with clades shaded and labelled by phylum and a branch-length scale bar. Three concentric rings surround the tree; orange: bar plot of the number predicted promoters per species, up to 10; green: GC content of the species; grey: genome lengths in bp. (B) Distribution of shortlist size across the species representatives, from none to the maximum of ten. Labels give the percentage and count of representatives returning no promoter and of those returning a complete shortlist. (C) Percentage of species representatives returning a complete shortlist against genome GC content, plotted at GC-decile medians and drawn separately within each tertile of genome size, with the size range of each tertile given. (D) Transfer of the marker panel across the domain, for the 26 most widely served gene symbols and the 28 phyla contributing most representatives. Cell colour is the percentage of that phylum’s representatives in which the gene is served by a shortlisted promoter, on a logarithmic scale; grey marks gene–phylum combinations never observed. (E) Use of the three spacings the motif search permits, expressed relative to equal use of all three; the dashed line is that equal-use baseline and the percentage of shortlisted promoters carrying each spacing is labelled. (F) Percentage of shortlisted promoters carrying a 17 bp spacer in each of the 67 phyla contributing at least 100 calls, ordered by that percentage, with exact binomial 95% confidence intervals. The dashed line is the equal-use baseline of one third; intervals are coloured by whether they exclude it. The phylum with the lowest value is labelled.

ProFinder returned a shortlist across the entire compositional range (Fig. 6C), with genomes between 39% and 65.3% GC returning a full ten promoters 87.3–92.6% of the time, compared to 70.6% at the GC-rich extreme and 48.4% at the AT-rich extreme (Table S44). Genomes at the AT-rich extreme were also the smallest (1.30 Mb compared to 4.05 Mb at the GC-rich extreme) and their shortfall disappeared once size was held constant, from 40.1% of small genomes returning a full ten to 96.4% of mid-sized and 100% of large ones (Fig. 6C). Thus, this was a consequence of genome size rather than of base composition. In contrast, the GC-rich decline persisted in all three size classes (90.5%, 53.8% and 13.9% of large, mid and small genomes; Fig. 6C), suggesting a direct GC effect. Nevertheless, the median genome at the GC-rich extreme retained 15 candidate promoters, more than a ProFinder shortlist reports, with only 29.4% of GC-rich genomes left with fewer than ten.

In total, 519,631 promoters were shortlisted, which were associated with 9,370 distinct genes (Table S45). As all genes within a defined operon qualified, these gene products were not restricted to the 98 profile families of the marker panel. A total of 29 genes were recovered from 50 or more phyla, led by the β sliding clamp *dnaN* in 95 phyla (8,996 promoters), RNA polymerase β subunit *rpoB* in 69 (8,659) and *recA* in 64 (7,718) (Fig. 6D, B; Table S46). Gene symbols that were detected only once accounted for 0.76% of calls (*n* = 3,140), while 115 symbols covered 50% of calls and 575 covered 80%, indicating that the shortlists converged on essential functions without collapsing onto a handful of genes (Fig. 6D).

The recovered promoters carried the canonical σ^70^ architecture. The three permitted spacings (16, 17, 18 bp) were not used equally, with 17 bp accounting for 43.6% of shortlisted promoters compared to 29.6% and 26.8% for 16 and 18 bp (χ^2^= 25,323, 2 d.f., *p* < 1e-300; Fig. 6E; Table S47). Of the 67 phyla with at least 100 calls, 17 bp was significantly enriched above the equal-use baseline in 60 and fell significantly below it in none, the remaining seven being indistinguishable from the baseline. The weakest was Caldisericota at 30.3% (95% CI 22.3–39.3%; Fig. 6F). This preference was unrelated to base composition across phyla (ρ = +0.07 compared to phylum mean GC, *p* = 0.57), so it was not an artefact of AT-rich hexamers in AT-rich genomes. The canonical spacer therefore holds across the bacterial domain.

Overall, ProFinder returns a ranked, marker-anchored promoter shortlist of similar length and gene content for the majority of bacterial species, wherever they sit in the phylogeny and whatever their genome size or base composition. For small or GC-rich genomes a reduced, yet usable shortlist is returned.

## Discussion

Characterised promoters are scarce in non-model bacteria, limiting heterologous expression and circuit construction in these organisms (7,35). The standard workaround is to repurpose intergenic regions (IGRs) upstream of housekeeping genes and screen candidates empirically (36). However, many of these candidates fail to drive transcription (37), multiplying the variables that require testing and optimisation during genetic tool development (36,38). Genome-wide promoter classifiers offer an alternative but show degraded accuracy on species phylogenetically distant from their *E. coli* and *B. subtilis* training data (39), and there is no indication whether the promoters are active (18). Phylogenomic marker gene panels (e.g., CheckM, Bac120/GTDB-Tk, AMPHORA2) identify single-copy housekeeping genes that are generally constitutively expressed (40–42), but they operate on coding sequences and provide no information about associated promoter elements or expression levels across conditions. In response, we developed ProFinder to address both gaps in one workflow.

Identifying conserved housekeeping genes alone is insufficient to guarantee strong, constitutive expression (43). The most direct confirmation would come from longitudinal transcriptional data across multiple conditions (44), yet such datasets remain unavailable for most microbes. As a practical proxy, we evaluated publicly available expression profiles measured across diverse environments and organisms (5). This analysis revealed that the number of conserved gene families exceeding a moderate expression floor drops off at a certain point (43), indicating that many otherwise conserved housekeeping genes are either too lowly expressed or too variable to serve as reliable constitutive parts. ProFinder enforces both broad expression across conditions and a minimum expression magnitude, distinguishing genes that are simply “not silent” from those actively maintained at levels suitable for biotechnological applications.

ProFinder predicts the promoters of these genes by exploiting an established biological prior: the majority of bacterial transcription start sites (TSSs) reside in the IGR directly upstream of the start codon, oriented toward the coding sequence (45,46). These criteria provide a sequence agnostic constraint that generalises across phylogeny. We further refine the search space according to established biophysical constraints on promoter length, ensuring sufficient room for the σ^70^ –35 and – 10 motifs plus the flanking contacts required by the RNA polymerase holoenzyme (47). An upper bound reflects the documented decline in TSS density in long IGRs, where functional promoter elements become increasingly diluted by non⍰regulatory sequence (45,46). This geometric window captures the interval most likely to harbour a σ^70^⍰dependent promoter while excluding extraneous DNA and substantially raising the signal to noise ratio of the subsequent motif search.

By design, these geometric criteria define a high-yield search space for promoters, but not an exhaustive one. Promoters located internally within long operons (45), regulatory sequences embedded in upstream coding regions (48), or the tightly packed promoters of extremely compact genomes are deliberately excluded. This design optimises for precision over recall, returning a short, high confidence shortlist of candidate promoters suited for genetic engineering rather than a complete atlas of all possible transcription initiation sites.

Within the filtered IGR set, ProFinder detects σ^70^promoters using a collection of PWMs built from diverse bacterial lineages. Although the –10 element is broadly conserved, the –35 hexamer exhibits considerable lineage⍰specific variation. A multi⍰organism PWM library enables ProFinder to capture this full spectrum while remaining anchored to the conserved ^70^ architecture.

Applying ProFinder to the entire bacterial domain tested whether these priors hold across cultured and uncultured diversity, including in phyla represented by a single genome. Almost every genome returned at least one promoter, and sparsely sampled lineages produced equivalent results to well-sampled ones so the method does not silently fail in unrepresented regions of sequence space. In AT-rich genomes, smaller genome sizes led to fewer predicted promoters, likely reflecting the limited marker complement of reduced genomes. In GC-rich genomes, by contrast, a compositional effect reduced the output. ProFinder’s dependence on composition is nonetheless weaker than that of single-organism classifiers and at both compositional extremes it returns a shorter shortlist rather than one less likely to be genuine.

Several limitations must be acknowledged. ProFinder’s reliance on a core set of housekeeping markers reduces predictions in heavily reduced genomes, such as those of obligate endosymbionts, that lack many of these orthologs (52,53). Furthermore, transcript stability, attenuation and translational control cannot be distinguished from initiation in these data, so the panel identifies genes whose transcription is maintained, not promoters whose output is unregulated. Comprehensive benchmarking remains restricted to *E. coli* and *B. subtilis*, the only two species with complete, experimentally validated promoter libraries. Although these organisms are phylogenetically distant (Proteobacteria and Firmicutes, respectively), they do not span the full bacterial domain. Nevertheless, this limitation mirrors the gap ProFinder is designed to address: most bacteria lack the genetic tools and datasets needed for systematic promoter characterisation. Until such resources become broadly available, ProFinder serves as a pre-computed, reliable starting point designed to accelerate this process.

In sum, ProFinder reframes promoter prediction for genetic engineering by layering three complementary biological priors: marker genes selected for conservation and for transcription maintained across conditions; CDS-oriented intergenic regions of empirically bounded length; and a multi-organism σ^70^ motif catalogue with its characteristic spacer. No layer identifies a promoter alone, and none is a trained promoter model, but applied together they reduce a genome to a short, ranked set of native candidates designed to accelerate the engineering of industrially relevant species, environmentally important microbes, and previously uncultivated lineages.

## Data Availability

The data underlying this article are available in the article and in its online supplementary material. The ProFinder source code can be accessed from: https://github.com/xxxxx (will be provided upon publication) and the webserver is freely available at: https://plabase.cs.uni-tuebingen.de/profinder/.

## Supplementary Tables

**Table S1. *Source marker gene sets used to construct the HMM profile collection.*** Ten published sets of conserved single-copy bacterial marker genes are listed. Each set is listed with its size (number of genes and associated HMM profiles), original reference, and a brief description of its intended application. The final row reports the union across all 10 sets.

**Table S2. *Gene membership across the 10 marker gene sets*.** A 194 × 10 binary matrix indicating the presence (1) or absence (0) of each gene in each source set. Genes are sorted by descending number of sets.

**Table S3. *Catalog of 838 HMM profiles.*** All HMM profiles.

**Table S4. *Pairwise Jaccard similarity among the 10 marker gene sets.*** The Jaccard indices computed over gene-level membership.

**Table S5. Gene families detected across all five reference organisms.** The 261 gene families for which hmmsearch identified a homolog in all five organisms (Escherichia coli MG1655, Bacillus subtilis 168, Mycobacterium tuberculosis H37Rv, Pseudomonas aeruginosa PAO1 and Salmonella enterica Typhimurium LT2), listed alphabetically with their member profiles and, for each organism, the matched locus tag and its full-sequence bit score.

**Table S6. *Per-organism expression statistics for all 261 gene families, annotated by filter outcome.*** For each family and each reference organism the table reports the matched locus tag, the number of conditions surveyed, and the median, mean and standard deviation of centred log_2_-TPM, together with the fraction of conditions above the −3 log_2_-TPM depletion threshold. Three cross-organism summaries follow: the expression floor (lowest of the five medians), the mean of the five medians, and the lowest of the five detection fractions. The filter outcome records whether a family failed the breadth filter, failed the magnitude filter, was removed as organism-sensitive after leave-one-out, or entered the final 98-family panel.

**Table S7. *Depletion threshold sweep.*** For each candidate depletion threshold from -8 to +2 log_2_- TPM, the number of gene families expressed above it in at least 95% of conditions in every organism, and the corresponding count within each organism separately.

**Table S8. *Leave-one-out robustness.*** Each row reports the result of excluding one reference organism and re-running the selection on the remaining four, giving the number of families passing each filter, the magnitude-filter cutoff, and the agreement with the full-panel result.

**Table S9. *Pairwise Jaccard similarity among leave-one-out panels.*** Jaccard indices between the family sets retained by the full run and by each leave-one-out run.

**Table S10. *The 98 marker gene families forming the final panel.***

**Table S11. *Per-family expression dynamic range*.** For each of the 261 conserved families, the span between its 2.5th and 97.5th expression percentiles in whichever organism that span is widest, given in log_2_ units and as a fold change, together with the interquartile range and standard deviation in that organism, the family’s median across organisms, whether it entered the 98-family panel, and its housekeeping class where it has one. Group medians quoted in the text are the median of the log_2_ spans converted to a fold change, which for an even-sized group is not the median of the fold-change column.

**Table S12. *Generalisation to six organisms, summary.*** Statistics comparing panel families with the equally conserved families the filters rejected, and with each genome’s own gene complement, across the six organisms not used to define the panel. Each row gives the quantity, its value and a note recording the sample size or the test applied.

**Table S13. *Generalisation to six organisms, per organism.*** For each of the six organisms, the median within-genome stability percentile of the panel families and the Kolmogorov-Smirnov *p*-value against the uniform null that a randomly chosen gene set would follow.

**Table S14. *Genome panel and phylum assignment.*** The 49 panel genomes with their taxonomic lineage.

**Table S15. *Per-genome transcription start site and intergenic region characteristics.*** One row per genome, giving intergenic region counts and total length, TSS counts by context, chromosome length and gene coverage, and the percentage of that genome’s TSSs lying in intergenic sequence. The TSS and intergenic counts summed over the panel are the figures reported in the text, and the intergenic and coding densities are computed from the total intergenic length and chromosome length given here.

**Table S16. *Intergenic regions across the 49 chromosomes.*** One row per region, giving its orientation class, length, and the number of TSSs it contains on each strand. Regions are delimited by all annotated gene-like features, not by coding sequences alone. Regions upstream of a start codon (USS) are the tandem+, tandem-and divergent classes only.

**Table S17. *Per-genome motif subgroups.*** The 127 subgroups recovered across the panel, each with its consensus −35 and −10 hexamers, its genome, and the number of promoters it contains.

**Table S18. *Motif architecture catalogue.*** The 72 distinct pairs of consensus −35 and −10 hexamers, with the number of promoters and genomes each accounts for and the cumulative share of catalogued promoters.

**Table S19. *Motif architectures per genome.***

**Table S20. *Partners per −10 hexamer.*** Each distinct consensus −10 hexamer with the number of distinct −35 hexamers it is catalogued with.

**Table S21. *Partners per −35 hexamer.*** Each distinct consensus −35 hexamer with the number of distinct −10 hexamers it is catalogued with.

**Table S22*. Share of promoters per architecture, separately for the −10 and −35 elements.***

**Table S23. *Partial Mantel correlations between hexamer usage and genome properties***. For each motif element, the correlation of between-genome hexamer usage distance with GC distance controlling for phylum, and with phylum controlling for GC, each with a permutation *p*-value.

**Table S24. *Distribution of the −35 / −10 spacer.*** Promoter counts and percentages at each spacer length observed in the catalogue.

**Table S25. *Per-genome distance from the −10 element to the start codon.*** For each panel genome, the number of promoters measured and the first quartile, median and third quartile of the distance.

**Table S26. *Distance-limit sweep.*** At each limit from 50 to 500 bp: the percentage of TSS-bearing intervals holding a catalogued promoter within the limit, pooled and as the lowest and highest values across genomes; the percentage in which a complete −10 / −35 pair was placed; and the percentage of calls that are TSS-bearing. Calls are scored leave-one-genome-out and reflect the motif gate alone.

**Table S27. *Per-genome reachability by distance limit.*** The percentage of each genome’s TSS-bearing intervals holding a catalogued promoter within each swept limit.

**Table S28. *Validation reference sets used for benchmark scoring.*** Two experimentally determined promoter sets were used to score predictions in *E. coli* K-12 MG1655 and *B. subtilis* subsp. subtilis 168. The *E. coli* reference is the σ^70^-annotated subset of RegulonDB v11 (10,738 promoters total). The *B. subtilis* reference is the union of σ^70^ (sigA) promoters annotated in DBTBS and SubtiWiki, comprising 4,005 promoters.

**Table S29. *Precision by tool in the two model organisms.*** For each of the five predictors in *E. coli* and *B. subtilis*, the number of calls scored, the number confirmed against that organism’s published promoter set, the resulting precision and its Wilson 95% interval. ProFinder is scored on all upstream-sense intergenic regions, with no shortlist cut-off and no marker-gene filter, so the comparison is of call quality rather than of the shortlist the tool is designed to produce.

**Table S30. *Top-ten shortlists for the model organisms.*** The ten ranked calls ProFinder reports for *E. coli* and *B. subtilis*, each with its interval, served gene and product, coordinates, the gating path that admitted it, its shortlist score, whether it matches the published reference set by homology and by coordinate overlap, the number of reference windows inside the interval and the distance to the nearest one.

**Table S31. *False positives among ten calls.*** For each model organism, ProFinder’s observed count in its ranked top ten with the gene and rank concerned and the resulting precision, against PromoterLCNN, which assigns no confidence score and so has no top ten: its value is the expectation over all draws of ten from its marker-gene sigma70 output, with the size of that pool and of its false-positive set, the 95% range, the modal count and its probability. The draw distribution is hypergeometric and enumerated exactly rather than simulated; the bootstrap mean is given alongside for comparison.

**Table S32. *Per-genome benchmark validation.*** One row per benchmark genome, sorted by GC content. Composition: the organism and strain, its GTDB phylum, GC content, the number of tandem intergenic regions supplied identically to both tools, whether the genome also contributed to the motif catalogue and under what name, and the first author, year and DOI of the study its promoter list comes from. Validation: the size of the published promoter list and how much of it resolved to coordinates, the proportion of annotated genes and of marker genes covered by that list, the number of marker-associated candidate promoters available, the number shortlisted and matched, and the composition-matched null rate with its binomial test.

**Table S33. *Overlap between the benchmark panel and the motif catalogue panel.*** For each benchmark genome, whether the organism also contributed to the motif catalogue, the matching catalogue entry, the basis of the match, and GC content.

**Table S34. *Benchmark validation across the 24-genome panel.*** For each genome, PromoterLCNN’s σ^70^ call count and expected validated count per ten random draws, ProFinder’s validated count among its ranked ten, and the size of its shortlist.

**Table S35. *Cross-species per-intergenic-region yield.*** For each of the 24 benchmark genomes, the number of intergenic regions, the ^70^ calls returned by ProFinder and by PromoterLCNN, the resulting per-region yields, and genome GC content and GC skew.

**Table S36. *RNA-seq runs for the extreme-GC organisms*.** One row per sequencing run used to quantify expression in the two highest-and two lowest-GC genomes of the benchmark panel, giving run, study and sample accessions, the organism, and the growth condition each run represents.

**Table S37. *Marker panel stability at compositional extremes.*** For the two highest-and two lowest-GC genomes, the number of RNA-seq runs and genes surveyed, how many panel families were detected and expressed in every condition, and the expression range of panel genes against other expressed genes.

**Table S38. *Shortlisted promoter expression at compositional extremes.*** For the same four genomes, the shortlist size, how many served genes were quantifiable and expressed in every condition, and the median expression of served genes against size-matched random draws from expressed genes with the associated *p*-values.

**Table S39. *Per-genome shortlist summary.*** One row per bacterial genome giving taxonomy, assembly quality, region and confirmed-call counts, shortlist size, calls by gating path, and genome size, GC content and contig count.

**Table S40. *Per-phylum summary.*** For each of the 149 phyla represented, the number of genomes, how many returned a complete or partial shortlist, the percentage partial, and median GC content and genome size.

**Table S41. *Shortlisted promoters detected across the 56,742 bacterial species.***

**Table S42. *Per-phylum enrichment of partial shortlists.*** For the 57 phyla with at least 20 genomes, the partial-return rate is tested against the panel-wide rate by exact binomial test, with Benjamini– Hochberg adjusted *p*-values.

**Table S43. *Per-phylum shortlist coverage.*** For each phylum, the number of genomes, the percentage returning a partial shortlist, and median genome size and assembly quality.

**Table S44. *Completeness by GC decile and size tertile.*** One row per decile of genome GC content, giving the decile bounds, the number of genomes, the percentage returning at least one promoter and the percentage returning a complete ten, that completeness broken down within each tertile of genome size, and the median shortlist length, candidate-pool size, motif-gate pass rate and genome size in the decile. The pooled and size-stratified completeness figures quoted in the text are the percent-complete column and the three tertile columns respectively.

**Table S45. *Served gene symbols by call count.*** Every gene symbol served by a shortlisted promoter across the bacterial representatives, with the number of calls it accounts for and the cumulative fraction of all calls down to that rank. Symbols are the annotation names carried by the genes the promoters serve, which is a larger and different set from the 98 profile families of the marker panel, because every gene of a qualifying operon is served.

**Table S46. *Marker occupancy by phylum.*** For the 26 most widely served marker genes, the percentage of each phylum’s genomes in which the gene is served by a shortlisted promoter, across the 28 best-sampled phyla.

**Table S47. *Spacer composition per phylum.*** For each phylum with at least 100 shortlisted promoters, the number of calls, the count and fraction at a 17 bp spacer with exact binomial 95% confidence limits, mean GC content, and whether the fraction lies above or below the equal-use baseline.

**Supplementary Figure 1.**
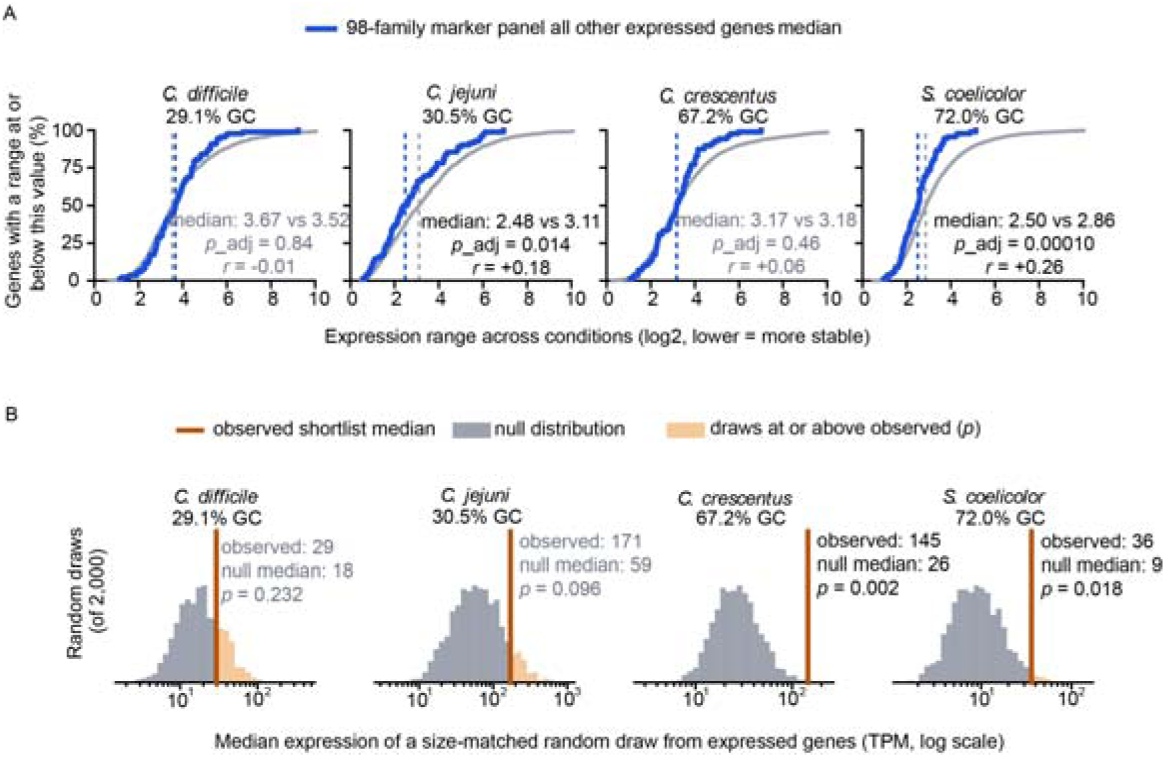
Marker-panel stability and shortlist expression at compositional extremes. Both panels cover the two lowest-and two highest-GC genomes of the 24-genome benchmark panel arranged left to right by increasing genome GC content, which is given below each organism name. Expression is quantified as TPM from public RNA-seq runs against a coding-sequence reference, and a gene counts as expressed where it reaches 1 TPM in at least half of that organism’s runs. (A) Cumulative distribution of each gene’s expression range across conditions, defined as the log2 ratio of the 97.5th to th 2.5th percentile of its TPM, so that a smaller value denotes a gene whose level varies less between conditions. Genes of the 98-family marker panel detected in that organism (blue) are plotted against all other expressed genes (grey); dashed vertical lines mark the two group medians. The annotation gives those medians, a two-sided Mann–Whitney U test of the two distributions with Benjamini–Hochberg adjustment across the four organisms, and a rank-biserial correlation r, which is positive where panel genes rank as the narrower set and negative where they do not; annotations are greyed and marked n.s. where the adjusted p does not reach 0.05. A curve lying to the left of the other denotes the more stable set at every quantile. (B) Null distribution of the median expression of a size-matched random draw from the same organism’s expressed genes. For each of 2,000 draws, k genes were taken without replacement and the median of their expression recorded, k being the number of genes served by that genome’s shortlisted promoters with quantifiable expression. The vertical line marks the observed median for the served genes and the shaded upper tail the draws at or above it; that fraction is the permutation p quoted, greyed where it does not reach 0.05. The horizontal axis is TPM on a logarithmic scale.

